# Fungi of the Fortuna Forest Reserve: Taxonomy and ecology with emphasis on ectomycorrhizal communities

**DOI:** 10.1101/2020.04.16.045724

**Authors:** Adriana Corrales, Clark L. Ovrebo

## Abstract

Panamanian montane forests harbor a high diversity of fungi, particularly of ectomycorrhizal (ECM) fungi, however their taxonomy and diversity patterns remain for the most part unexplored. Here we present state of the art fungal taxonomy and diversity patterns at Fortuna Forest Reserve based on morphological and molecular identification of over 1,000 fruiting body collections of macromycetes made over a period of five years. We compare these new results with previously published work based on environmental sampling of *Oreomunnea mexicana* root tips. We compiled a preliminary list of species and report 22 new genera and 29 new fungal species for Panama. Based on fruiting body collection data we compare the species composition of ECM fungal communities associated with *Oreomunnea* stands across sites differing in soil fertility and amount of rainfall. We also examine the effect of a long-term nitrogen addition treatment on the fruiting body production of ECM fungi. Finally, we discuss the biogeographic importance of Panama collections which fill in the knowledge gap of ECM fungal records between Costa Rica and Colombia. Given that the isthmus of Panama was an important migration route of ECM tree and fungal species from northern temperate areas to South America, the ECM fungal communities of Panama might show a high degree of isolation and therefore a high level of endemism. We expect that the forests at Fortuna will continue to yield new ECM macromycete data as we continue to study the collected specimens and describe new species.

## INTRODUCTION

Fungi play essential roles in tropical ecosystems. They are involved in organic matter decomposition, plant nutrition, nutrient cycling, and in some cases can regulate the local abundances of plants (Tedersoo et al., 2014). In neotropical montane cloud forests fungi are highly diverse and show strong endemism patterns (Del Olmo-Ruiz et al., 2017). In a recent review of fungal species associated with tropical montane cloud forests, Del Olmo-Ruiz et al. (2017) found 2,962 fungal species distributed within the neotropics. Of that total, only 220 species were originally described from neotropical montane forests while many others species were shared mostly with the northern temperate region (Del Olmo-Ruiz et al., 2017). This study suggests a large gap in the knowledge of fungal species from underexplored areas in Central and South America where a great part of their diversity likely remains unknown. In the case of Panama, Del Olmo-Ruiz et al. (2017) only report 23 species of fungi from montane cloud forests. Even though Panama has a detailed checklist of fungi containing records of 2,772 species for the country (based on Piepenbring, 2007, 2006, maintained at http://biogeodb.stri.si.edu/fungi), most of the species are from lowland forests and therefore there is a lack of knowledge of fungal communities from Panamanian montane forests.

In forested ecosystems, saprotrophic and mycorrhizal fungi are the most abundant functional groups of fungi. Saprotrophic fungi decompose organic matter into more simple carbohydrates that can be used for energy. Mycorrhizal fungi form symbiotic associations with the roots of plants receiving sugar from their host while helping them to obtain nutrients from the soil. Mycorrhizal fungi can be classified into two main groups: ectomycorrhizal (ECM) and arbuscular mycorrhizal (AM). Tropical montane forests between 1500– 2500 m a.s.l. are particularly rich in ectomycorrhizal (ECM) fungal species due to high density of ECM host trees, intermediate temperatures, and high precipitation (Corrales et al., 2018; Geml et al., 2014; Gómez-Hernández et al., 2012). The diversity patterns of fungi associated with ECM host plant species have been studied on only a few occasions in montane forests in tropical America. Fungi associated with *Quercus* dominated forests in Mexico, Costa Rica, and Colombia are the best known montane ECM systems in the neotropics (Franco-Molano et al., 2010; Halling and Mueller, 2002, 1999; Halling and Ovrebo, 1987; Morris et al., 2008; Mueller and Halling, 1995) starting with the pioneering work of Singer (1963) in oak forest of Colombia. *Alnus* dominated forests in Mexico, Bolivia, and Argentina have been studied to a lesser extent, for the most part, based upon sequencing of root symbiotic fungi using metagenomic approaches (Kennedy et al., 2015, 2011; Põlme et al., 2013; Wicaksono et al., 2017).

This chapter summarizes the research done at the Fortuna Forest Reserve (Fortuna) on the taxonomy and ecology of macrofungi. For the most part, we describe the communities associated with the ECM host tree *Oreomunnea mexicana* (Juglandaceae) based on new data from fruiting body surveys and sequencing of ECM colonized root tips previously published by Corrales et al. (2016, 2017). Other ECM host plants, including *Quercus* spp., *Coccoloba* spp., and *Alfaroa costaricensis* are also present in the Fortuna at lower abundances and therefore some of the species included in this chapter may be shared or associated with these species as well. To our knowledge this is the first study that uses fruiting bodies to characterize the fungal communities associated with *Oreomunnea mexicana*. This work, which focuses on the morphological and molecular identification of voucher specimens, not only constitutes an essential tool to document molecular ecology studies at Fortuna Forest Reserve but also provides a resource for other studies focusing on the ecology and biogeography of macrofungi in tropical montane forests in Central America and northern South America.

## METHODS

Fruiting body collections of macromycetes were made mainly in stands of *Oreomunnea mexicana* located in four watersheds in the Fortuna Forest Reserve (Fortuna): Alto frio (AF), Hornito (HO), Honda (HA and HB), and Zarceadero (Za) (for a detailed site description of AF, HO, HA and HB see Corrales et al. 2016a). Two different studies involving collection of fruiting bodies were done at these sites. First, during February to July 2012 we established 50 × 4 m transects that were revisited every 2 weeks to collect fruiting bodies of all the species of macromycetes present. This study aimed to compare mushroom diversity in sites with contrasting soil fertility and rainfall conditions. Second, during September 2013 to January 2014 collections were also made in a nitrogen (N) addition experiment that has been running since 2006 in the Honda watershed (Corre et al., 2010). Transects of 40 × 4 m were established in three N addition and four control plots containing *Oreomunnea* trees. These transects were revisited every 2 weeks with the aim of determining the effect of N addition on the production of ECM fruiting bodies.

To augment the data from the transects, we collected fruiting bodies opportunistically in areas near the transects and other sites at Fortuna where *Oreomunnea mexicana* is present along with *Quercus* spp. These collections were made over five years, between 2011 and 2015. The intent of this collecting effort was to capture diversity that might have been missed along the transects in order to ensure a more complete inventory.

Macromorphology of fresh fruiting bodies was recorded in the field, and a tissue sample was preserved for DNA extraction. Vouchers of fruiting bodies are deposited at the University of Arizona Robert L. Gilbertson Mycological Herbarium (MYCO-ARIZ), the herbarium of University of Central Oklahoma (CSU), and Herbario de la Universidad de Panama (PMA). Genomic DNA was extracted using the REDExtract-N-Amp tissue PCR kit (following the manufacturer’s instructions; Sigma-Aldrich). Primers ITS1F, ITS4, and ITS4B were used for DNA amplification. Methods for species molecular identification follows (Corrales et al., 2016). Voucher collections were identified based on their morphology and DNA with the help of specialists in the group and names assigned when possible. Because of the high diversity and challenging taxonomy of many of the groups, and the possibility that collections represent new species, some species names as well as total diversity of certain genera were assigned based on sequence similarity with a 97% threshold used to classify fungal operational taxonomic units (OTUs).

### PRELIMINARY LIST OF FUNGAL SPECIES FOR THE FORTUNA FOREST RESERVE WITH NEW REPORTS FOR PANAMA

During five years of collection, a total of 1,003 fruiting body collections were made and processed as herbarium specimens. In addition, 400 of these collections were sequenced to obtain fungal barcode data (ITS region). This section summarizes the most up-to-date results that we have for the fungal inventory. The list of fungi of Fortuna is still a work in progress given that many of the collected specimens likely represent new species. Currently we are in the process of describing more than 20 species in collaboration with the specialists in each taxonomic group.

We have found a total 157 species (including morphospecies and OTUs), with most species belonging to Basidiomycota and only 4 species belonging to Ascomycota. These species are distributed in 65 genera (28 ECM, 35 saprotrophs, and 2 fungal parasites) and 35 families (14 ECM, 20 saprotrophs, and one fungal parasite; Table 1). The most species-rich genus was *Russula* with about 40 species (defined based on OTUs, Corrales et al. *in prep*) and *Lactarius* with 8 species. The family with the most genera was Boletaceae with 11 genera followed by Russulaceae, Entolomataceae, Hymenogastraceae, Marasmiaceae, Mycenaceae, Phallaceae, and Physalacriaceae all of them with 3 genera (Table 1).

**Table 1.** Estimates of fungal species per family and per genus based on morphology and DNA of fruiting body surveys at Fortuna. ** New genus records for Panama, ^1^ Presumably biotrophic genus

| Taxa | No. species | Taxa | No. species | Taxa | No. species |
| --- | --- | --- | --- | --- | --- |
| **Ectomycorrhizal** | **48** | **Ectomycorrhizal (continued)** |  | **Saprotrophic (continued)** |  |
| **Amanitaceae** | **8** | **Leotiaceae** | **2** | **Meruliaceae** | **2** |
| *Amanita* | 8 | *Leotia* | 2 | *Cymatoderma* | 1 |
| **Bankeraceae** | **2** | **Russulaceae** | **51** | *Podoscypha* | 1 |
| *Hydnellum* ** | 1 | *Lactarius* | 8 | **Mycenaceae** | **4** |
| *Phellodon* | 1 | *Lactifluus* ** | 3 | *Filoboletus* | 1 |
| **Boletaceae** | **20** | *Russula* | 40 | *Mycena* | 2 |
| *Austroboletus* ** | 1 | **Sclerodermataceae** | **1** | *Xeromphalina* | 1 |
| *Aureoboletus* ** | 1 | *Veligaster* | 1 | **Omphalotaceae** | **4** |
| *Boletellus* | 3 | **Saprotrophic** |  | *Gymnopus* | 3 |
| *Boletus* ** | 1 | **Agaricaceae** | **2** | *Marasmiellus* | 1 |
| *Leccinum* | 2 | *Agaricus* | 1 | **Phallaceae** | **3** |
| *Phylloporus* | 6 | *Leucocoprinus* | 1 | *Aseroë* ** | 1 |
| *Retiboletus* ** | 1 | **Auriculariaceae** | **1** | *Laternea* | 1 |
| *Strobilomyces* ** | 1 | *Auricularia* | 1 | *Phallus* | 1 |
| *Tylopilus* ** | 3 | **Auriscalpiaceae** | **1** | **Psathyrellaceae** | **1** |
| *Veloporphyrellus* ** | 1 | *Artomyces* ** | 1 | *Psathyrella* | 1 |
| **Calostomataceae** | **1** | **Entolomataceae** | **4** | **Physalacriaceae** | **3** |
| *Calostoma* | 1 | *Alboleptonia* | 1 | *Armillaria* | 1 |
| **Cantharellaceae** | **4** | *Inocephalus* ** | 1 | *Cyptotrama* ** | 1 |
| *Cantharellus* | 2 | *Entoloma* * | 2 | *Xerula* | 1 |
| *Craterellus* | 2 | **Fistulinaceae** | **1** | **Polyporaceae** | **2** |
| **Cortinariaceae** | **11** | *Fistulina* | 1 | *Lentinus* | 1 |
| *Cortinarius* | 8 | **Ganodermataceae** | **1** | *Polyporus* | 1 |
| *Inocybe* ** | 3 | *Ganoderma* | 1 | **Strophariaceae** | **1** |
| **Elaphomycetaceae** | **1** | **Hygrophoraceae** | **3** | *Agrocybe* ** | 1 |
| *Elaphomyces* ** | 1 | *Hygrocybe* | 3 | **Tapinellaceae** | **1** |
| **Gomphaceae** | **1** | **Hymenochaetaceae** | **1** | *Tapinella* ** | 1 |
| *Ramaria* | 1 | *Hymenochaete* | 1 | **Tricholomataceae** | **1** |
| **Hydnaceae** | **2** | **Hymenogastraceae** | **3** | *Clitocybe* | 1 |
| *Hydnum* ** | 2 | *Gymnopilus* | 1 | **Fungal parasite** |  |
| **Hydnangiaceae** | **4** | *Psilocybe* | 1 | **Boletaceae** | **1** |
| *Laccaria* | 4 | **Marasmiaceae** | **8** | *Chalciporus\*\** | 1 |
| **Hymenogastraceae** | **1** | *Crinipellis\*\** | 1 | **Ophiocordycipitaceae** | **1** |
| *Phaeocollybia\*\** | 1 | *Gerronema* | 1 | *Tolypocladium\*\** | 1 |
|  |  | *Marasmius* | 6 |  |  |

After revising the checklist of Panamanian fungi (http://biogeodb.stri.si.edu/fungi) and recent mycological publications from Panama, we report for the first time 22 new genera for the country. For ectomycorrhizal fungi, 13 out of 28 genera were new records for the country while for saprotrophic fungi, 7 out of 36 genera were new records. Both genera of fungal parasitic fungi were also new records for Panama. From the specimens that were possible to identify to species level, we report 29 new fungal species records for Panama (17 ECM and 12 saprotrophs) (Table S1). This is not surprising given that the checklist maintained on the STRI website includes very few ECM genera among the 657 species of Agaricomycetes reported from Panama. Finally, while 12 out of the 26 ECM basidiomycete genera listed in Table 1 are on the STRI checklist, they mostly indicate the presence of only a single species. Exceptions are *Russula* with seven species and *Ramaria* with six species listed.

### GENERIC DIVERSITY PATTERNS OF ECTOMYCORRHIZAL FUNGI ASSOCIATED WITH STANDS OF *OREOMUNNEA MEXICANA* GROWING ON SITES WITH CONTRASTING ABIOTIC CONDITIONS

Based on 297 fruiting body collections, made in sites with contrasting soil fertility and rainfall conditions, we found that overall the most abundant genera at Fortuna were *Russula*, *Lactarius*, *Laccaria*, *Cortinarius*, *Cantharellus*, *Boletus*, and *Amanita* (Appendix: Plates 1-4). At the genus level, HA and AF sites show the highest number of genera with 19 and 18 genera respectively. From the seven most abundant ECM genera only *Amanita*, *Laccaria*, and *Russula* were found across all sites while *Cortinarius* (not collected in AF and ZA), *Lactarius* (not collected in ZA), *Boletus* (not collected in HB), and *Cantharellus* (not found in HO) were found in some sites but not in all (Figure 1). Interestingly, few of the genera missing from some transects were later collected during the opportunistic surveys. It could be that some of these taxa are only symbionts of *Quercus,* or other ECM hosts present in low abundance at these sites, or that they produced fruiting bodies at times when collecting was not done.

**Figure 1.**
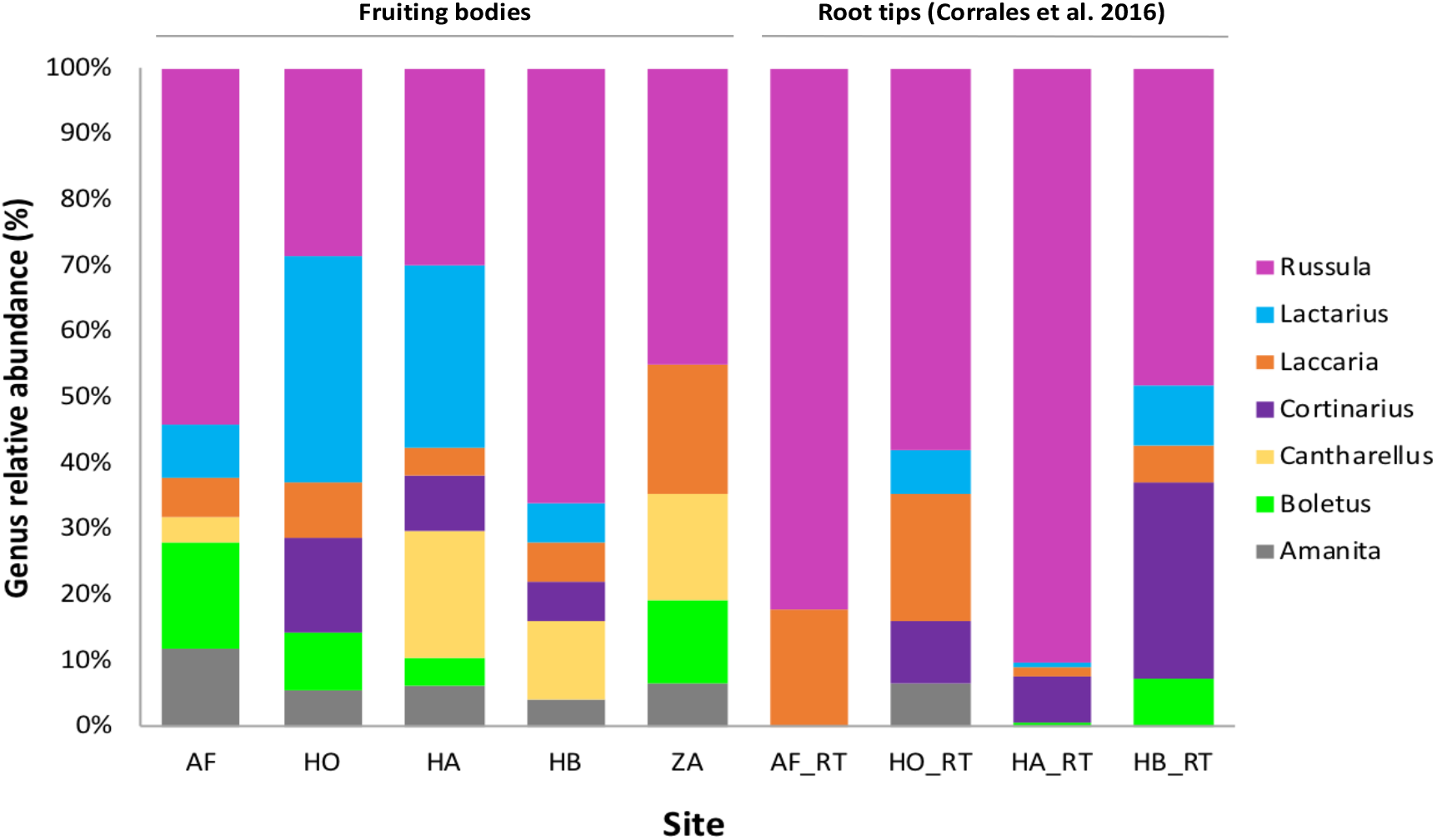
Number of collections for the seven most abundant ECM genera in each of the sites sampled along transects at Fortuna using fruiting bodies (left) and root tips (right; using data published by Corrales et al. 2016). Site abbreviations are: Alto Frio (AF), Hornito (HO), Honda A (HA), Honda B (HB), and Zarceadero (ZA). Root tip (RT).

Previous molecular inventories using *Oreomunnea mexicana* root tips have found that fungal ECM communities at Fortuna show high species richness and compositional turnover among sites with contrasting soil and rainfall conditions (Corrales et al. 2016). Given that most of our fruiting body collections were made at same transects studied by Corrales et al. (2016) it allows a direct comparison of the results from fruiting body collections (above ground surveys) with root tips inventories (below ground surveys). Overall, *Russula*, *Cortinarius*, *Laccaria*, and *Lactarius* were found among the top seven most abundant genera for both above and below ground surveys (Figure 1). However, *Tomentella*, *Byssocorticium*, and *Elaphomyces* were either rare or absent from fruiting body surveys while these genera round out the top seven most abundant genera in root tip inventories. This is an expected finding given that these genera produce inconspicuous or truffle like fruiting bodies that are usually overlooked in above ground inventories (Koljalg et al., 2000; Tedersoo et al., 2014). However, *Cantharellus*, the third most abundant genus in terms of number of fruiting body collections, was also absent from root tip inventories. This could be due to the very long and variable ITS region of *Cantharellus* that is usually difficult to amplify using universal fungal primers ITS1F/ITS4 (Buyck et al., 2014).

Based on rarefaction curves constructed using 167 fruiting body collections that have been sequenced and assigned to OTUs (Figure 2), we found that the site with the strongest seasonality in moisture availability, AF, had the highest alpha diversity of ECM fungal species while the continually moist site, HB, had the lowest. This contrasts with the results from belowground surveys where AF showed the lowest alpha diversity of OTUs (Corrales et al., 2016). This pattern could be associated with a higher investment in dispersal or reproduction of species growing on sites that are at least temporarily limited by water in contrast with a higher investment on mycelium growth in species that are not limited by this resource (Ekblad et al., 2013).

**Figure 2.**
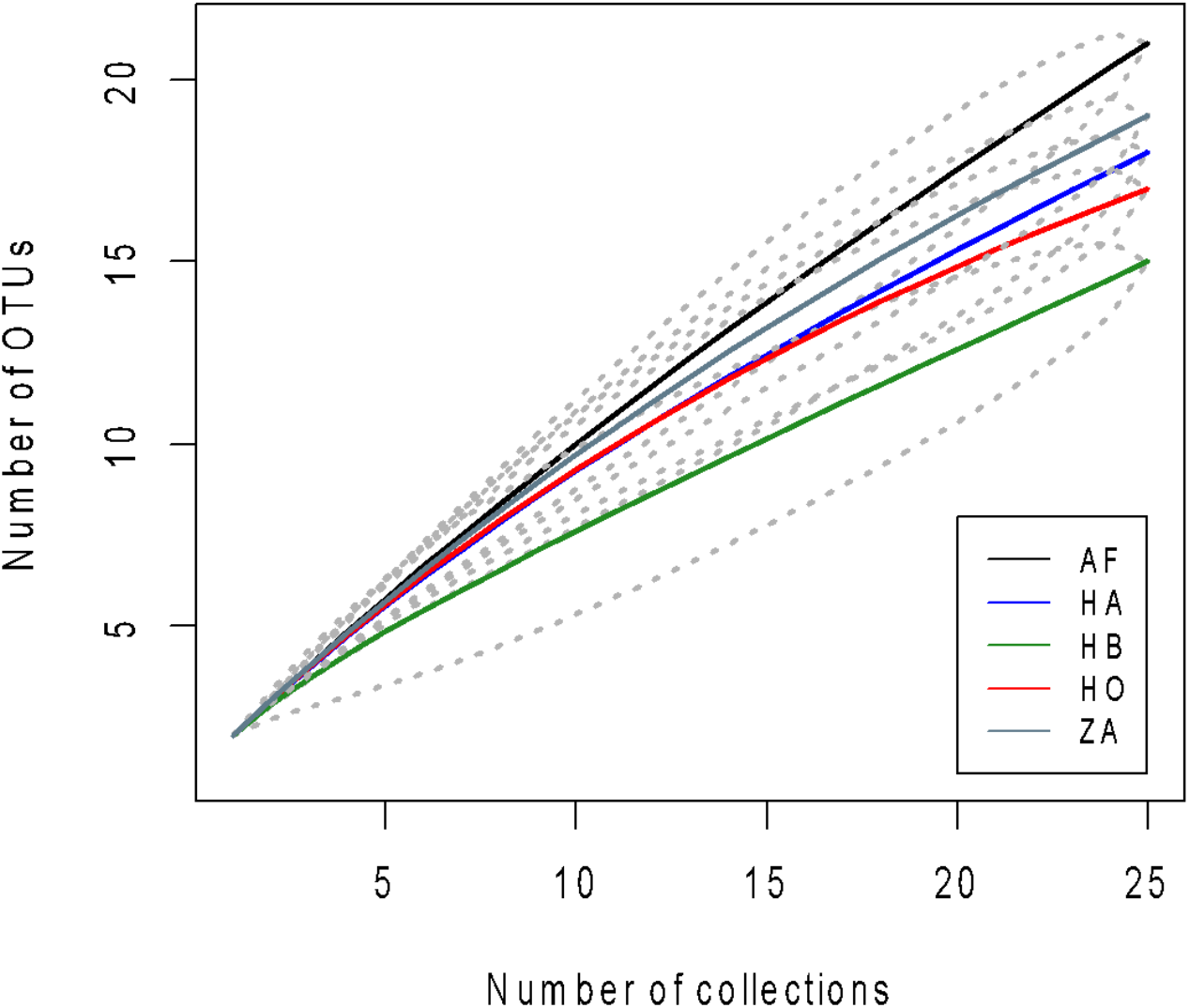
Ectomycorrhizal species accumulation curves and 95 % confidence per site based on OTUs from fruiting bodies. Site abbreviations are: Alto Frio (AF), Hornito (HO), Honda A (HA), Honda B (HB), and Zarceadero (ZA).

The timing of fruiting body production, or mushroom phenology, seems to be heavily influenced by differences in the rainfall patterns among sites at Fortuna. Based on fruiting body collections and rainfall data from 2012 (Dalling et al., *unpublished*), sites with a marked dry season between December and April (HO and AF) showed a strong peak of fruiting body production at the beginning of the rainy season between late April and June, similar to phenological observations obtained in nearby lowland vegetation (Piepenbring et al., 2015). This was most noticeable at AF, where more than 50 species were found fruiting in just one week (Figure 3A). In sites where the total amount of rainfall is higher and more evenly distributed throughout the year (HA and HB), fruiting body production occurred over the entire collecting period with peaks of fructification in February, April, and June (Figure 3B). Even though this is a small window of time to make generalizations about the local mushroom phenology, this is a first approximation to an aspect of fungal ecology that has been little studied in tropical forests. Over the years, we have also observed that the timing of the rainy season and the peak of fruiting body production could be very variable and is becoming more unpredictable due to local climate change. In future research, it would be interesting to monitor these changes over a longer period of time to understand the dynamics of the reproductive cycles of ECM fungi and the potential implications that climate change could have on those dynamics at Fortuna.

**Figure 3.**
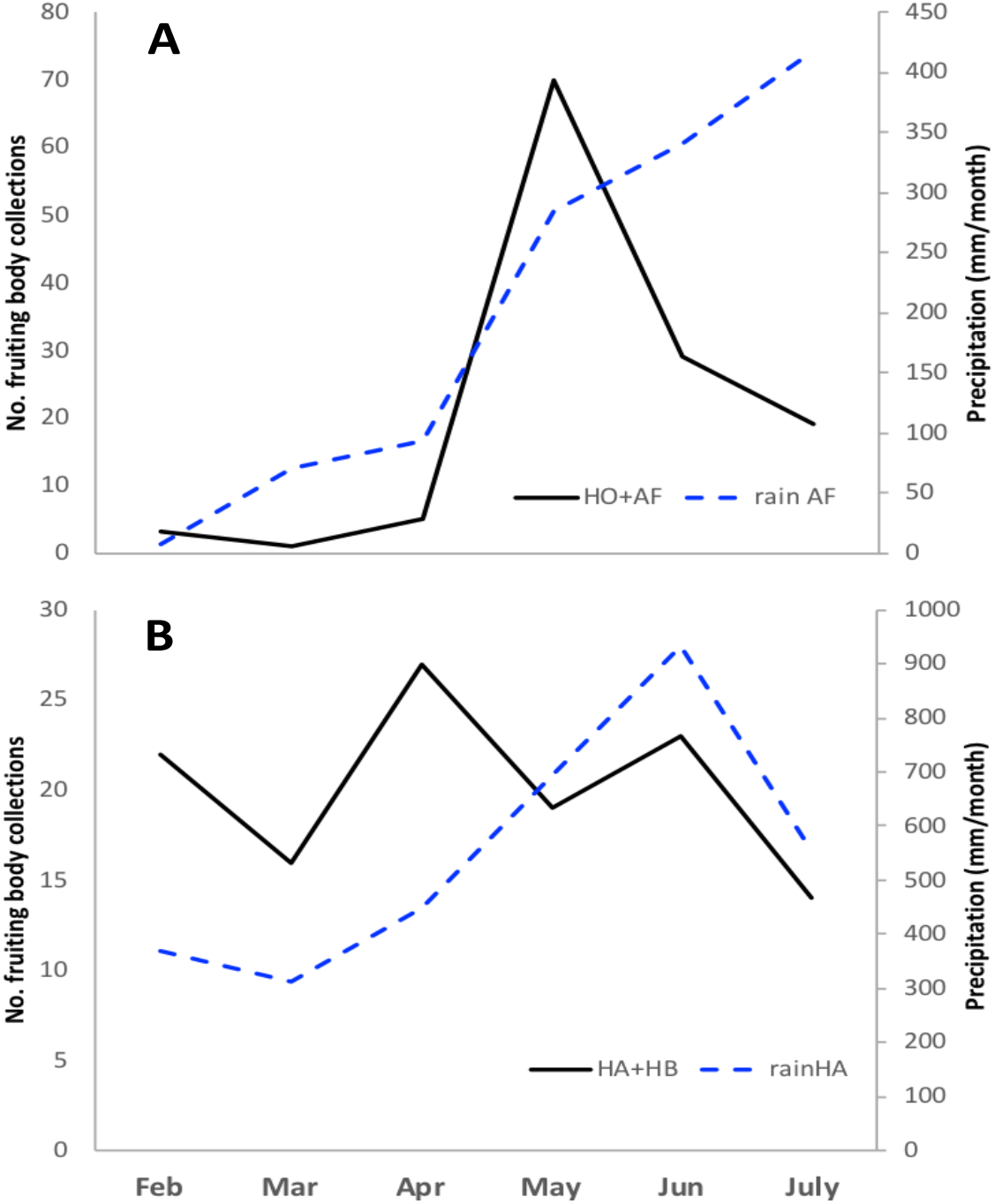
Number of fruiting body collections (solid line) during February to July 2012 and the local precipitation recorded at each site (dashed line). Number of collections were grouped by sites with differences in rainfall patterns following Corrales (2016a) **A**. Sites with dry period with < 100 mm of rainfall (AF: Alto Frio and HO: Hornito) and **B**. Sites with no months with < 100 mm of rainfall (HA: Honda A and HB: Honda B).

### INFLUENCE OF NITROGEN ADDITION ON ECTOMYCORRHIZAL FRUITING BODY PRODUCTION, DIVERSITY, AND SPECIES COMPOSITION

It is well recognized that ECM fungal communities respond to increases in N availability associated with anthropogenic fertilization or N deposition for both above and below ground structures (Lilleskov et al., 2011; van der Linde et al., 2018). However, most of these studies has been done in temperate or boreal forest and information about tropical ECM communities is scarce or nonexistent. Results from environmental sequencing of *Oreomunnea mexicana* root tips at Fortuna suggest that N fertilization could be associated with changes in ECM fungal community composition and could cause a reduction in soil enzyme activity and ECM root colonization (Corrales et al., 2017).

Surveys of ECM fruiting bodies done along transects at the same experimental plots used by Corrales et al. (2017), show that N addition could also have an effect on the overall production of fruiting bodies and frequency of fructification of some ECM genera. A total of 313 mushroom collections were made along seven transects. In control plots, we made an average of 44 (SD= 9) collections per plot belonging to 15 (SD= 1) genera while in N addition plots we made an average of 46 (SD= 4) collections per plot belonging to 12 (SD= 0.6) genera. This lower generic diversity found in N addition plots was also observed in the species-accumulation curves based on OTUs (obtained from sequencing of fruiting body collections) that show a lower species alpha diversity of the mushroom community present in N addition plots compared with control plots (Figure 4A). Community composition, however, did not show significant differences based on ordination and Adonis analyses performed using mushroom species presence-absence (Figure 4B) and abundance data (not shown) contrasting with results found by Corrales et al. (2017) based on environmental sequencing of *Oreomunnea* root tips that showed a significant change in species composition based on NMDS and Adonis analysis.

**Figure 4.**
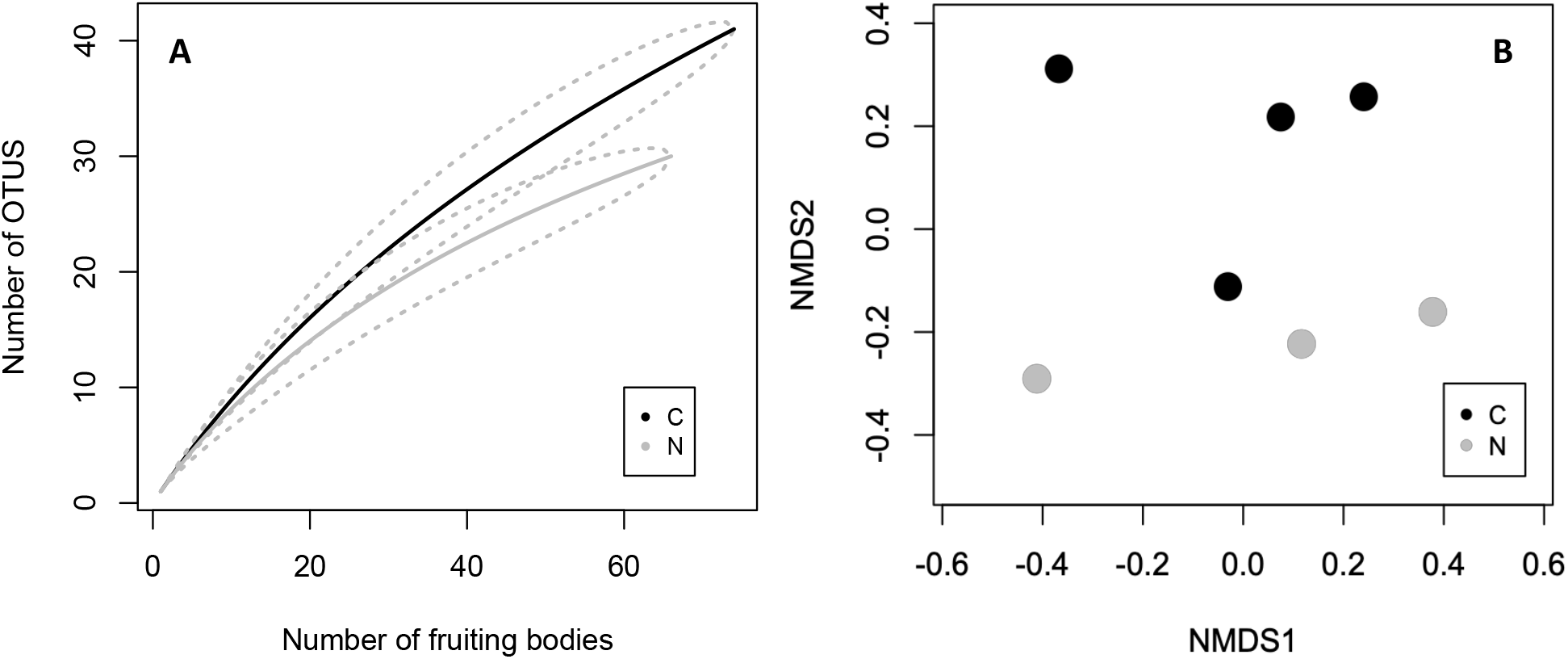
**A**. Species-accumulation curves of ECM fungi for N treatments (C: control and N: nitrogen addition) based on OTUs obtained from sequencing of fruiting body collections, **B**. NMDS for the mushroom community from control and nitrogen addition plots based on presence absence data and excluding singletons. Adonis analysis (P(>F)= 0.23, R^2^= 0.244).

*Russula, Lactarius, Cortinarius,* and *Leotia* were the most abundant genera in plots from both treatments (Figure 5) accounting for about 50% of all the collections. *Russula* and *Cortinarius* showed a strong reduction in the number of fruiting body collections between control and N addition treatments. A total of 41 collections of *Russula* were made in the control plots versus only 17 collections made in the N addition plots. For *Cortinarius* there were 18 collections made in control plots and 8 in N addition plots. In contrast, *Leotia* showed a much higher number of fruiting body collections in N addition compared with control plots with 17 and 10 collections respectively (Figure 5). Results based on environmental sequences of *Oreomunnea* ECM root tips published by Corrales et al (2017) also showed a lower number of sequences of *Cortinarius* in N addition plots compared with control plots but showed a higher abundance of *Laccaria* and *Lactarius* in N addition plots compared with control plots that was not reflected in the production of fruiting bodies.

**Figure 5.**
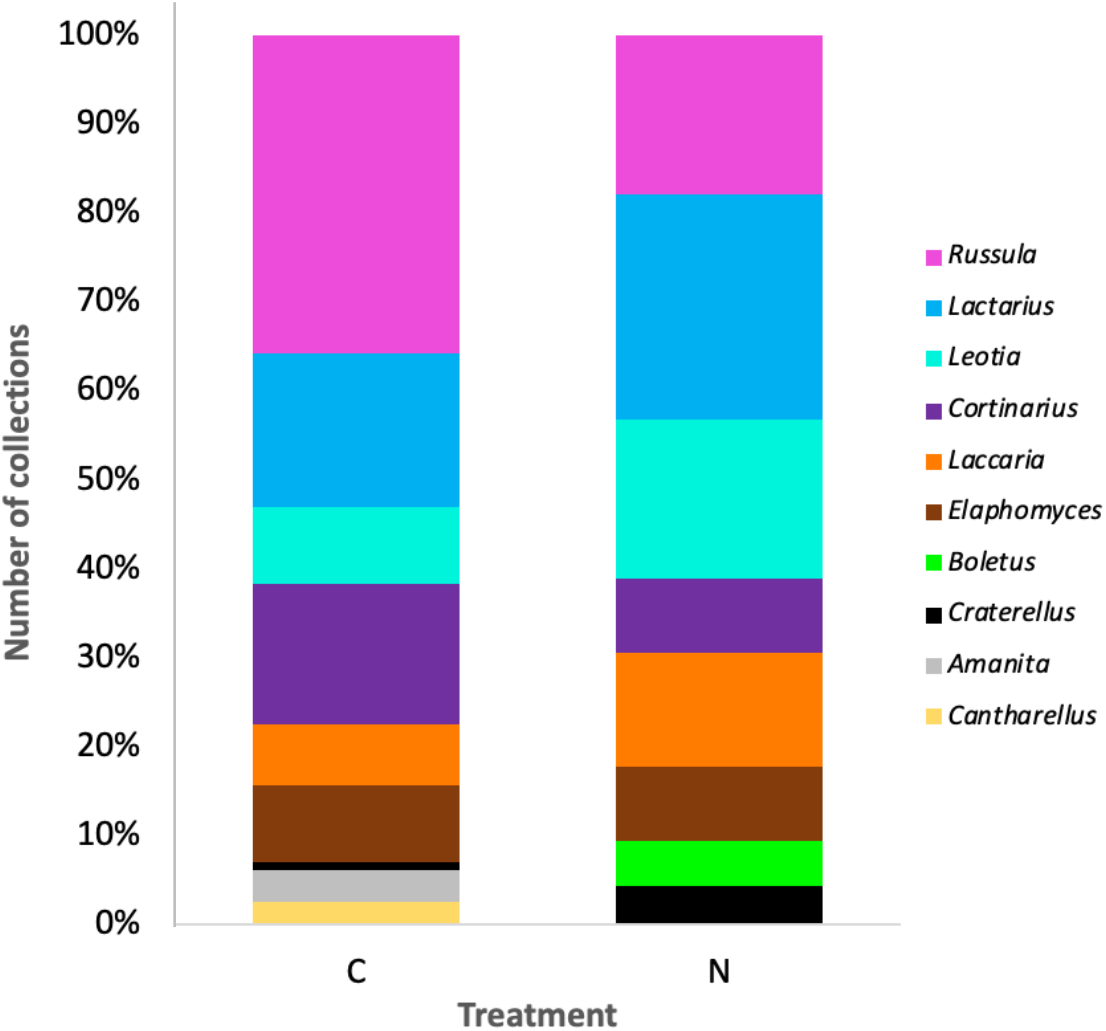
Number of fruiting body collections for the most abundant ECM genera in control (C) and nitrogen (N) addition plots sampled using transects at Fortuna.

**Figure 7.**
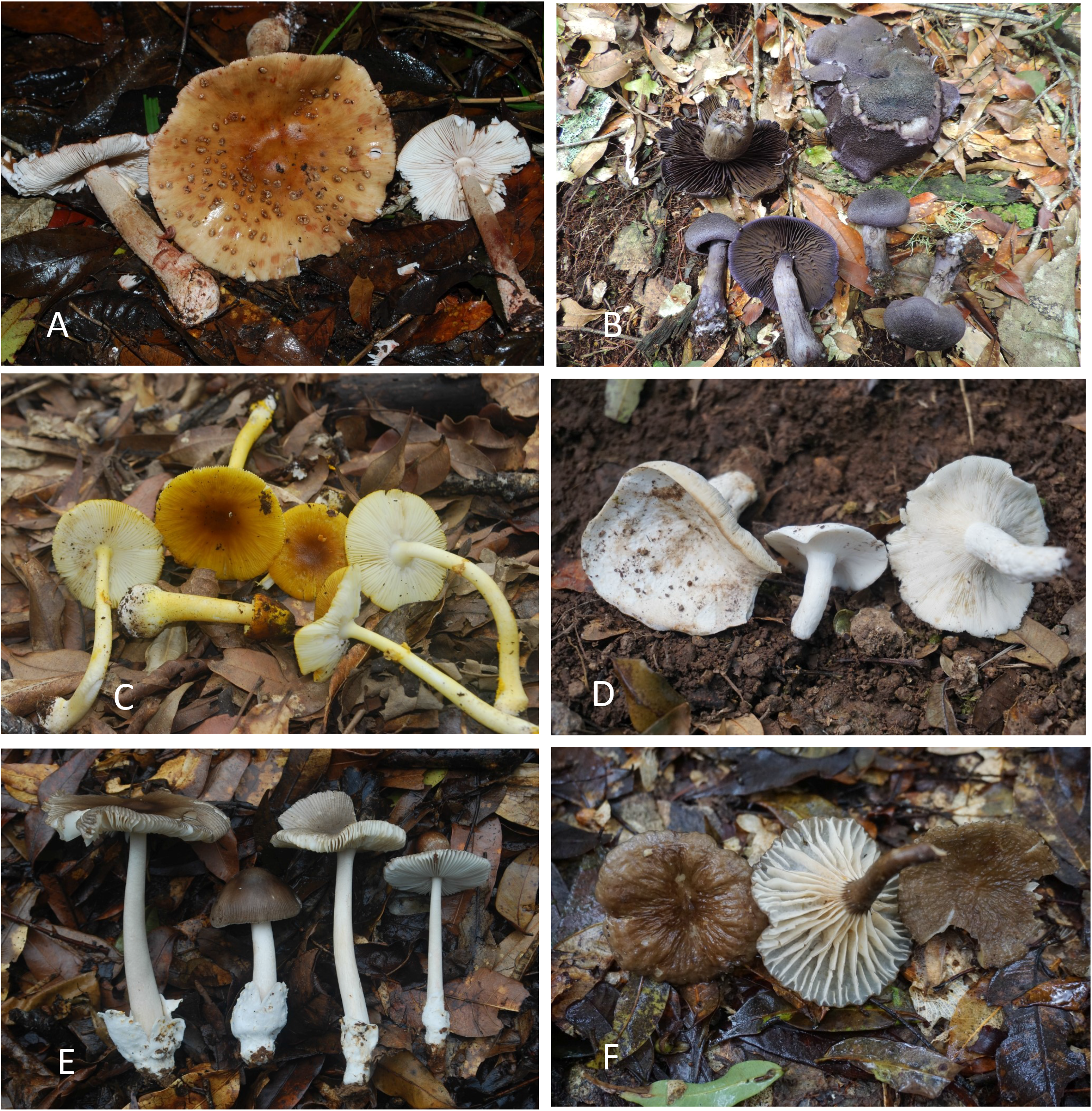
A-F. A. *Amanita brunneolocularis*. B. *Cortinarius neotropicus*. C. *Amanita flavoconia* var. *in-quinita*. D. *Lactarius* aff. *piperatus*, E. *Amanita* sp. F. *Lactarius* sp.

**Figures 8.**
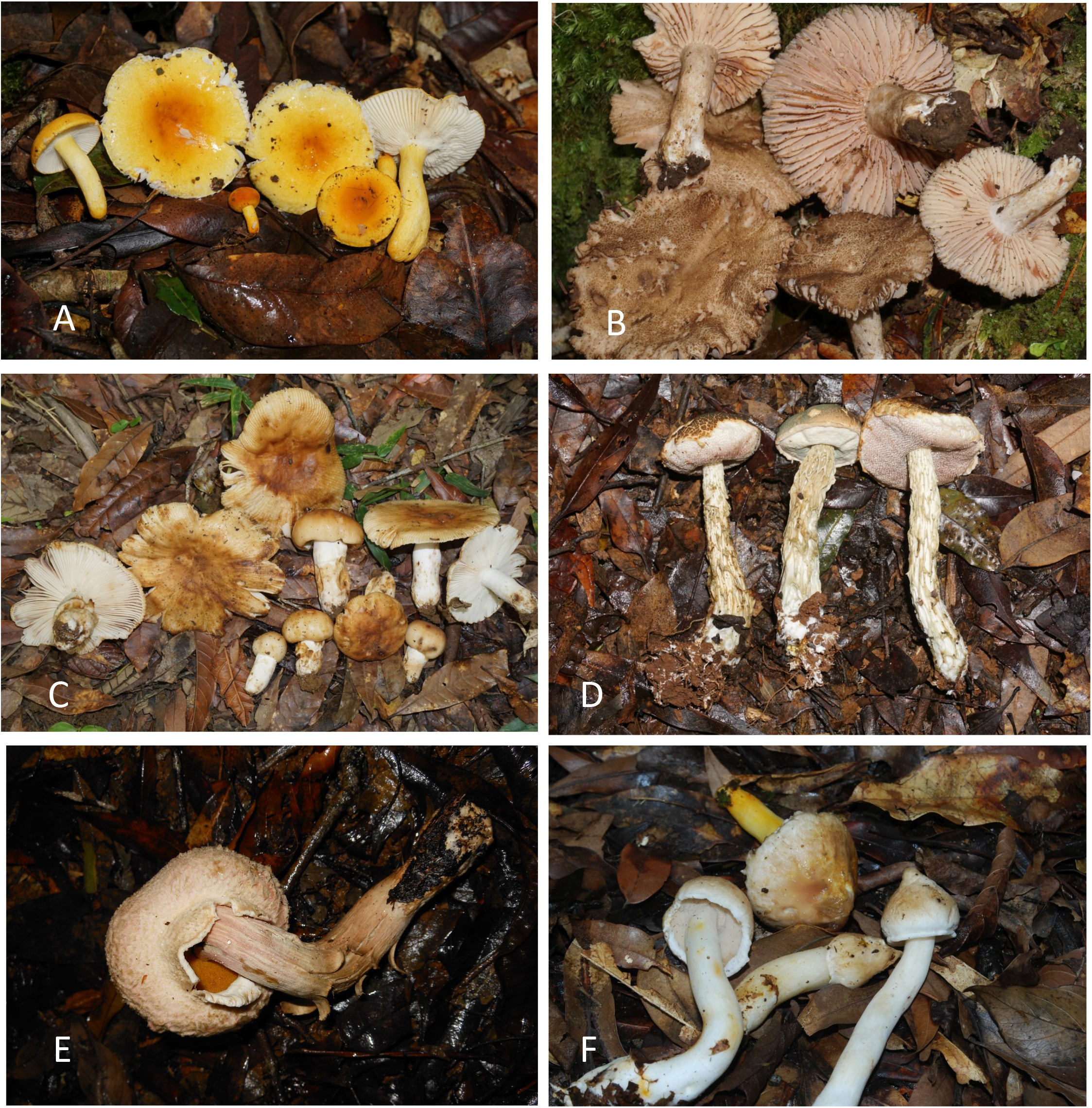
A-F. A. *Russula* sp. B. *Russula* sp. C. *Russula* sp. D. *Austroboletus neotropicalis*. E. *Boletellus ananas*. F. *Veloporphyrellus pantoleucus*.

**Figures 9.**
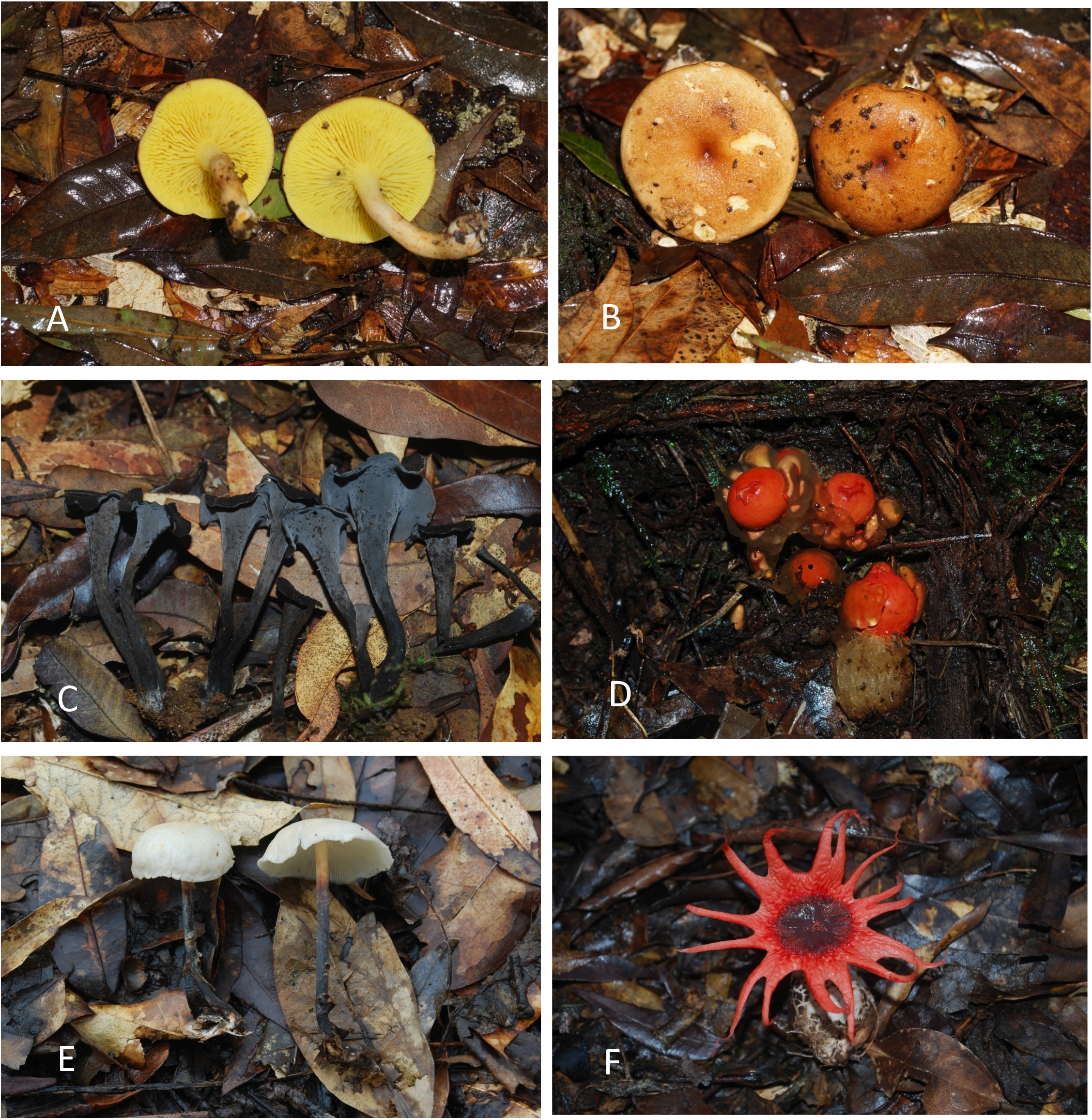
A-F. A-B. *Phylloporus centroamericanus*. C. *Craterellus* sp. D. *Calostoma cinnabarina.* E. *Gymnopus omphalodes*. F. *Aseroë rubra*.

**Figures 10.**
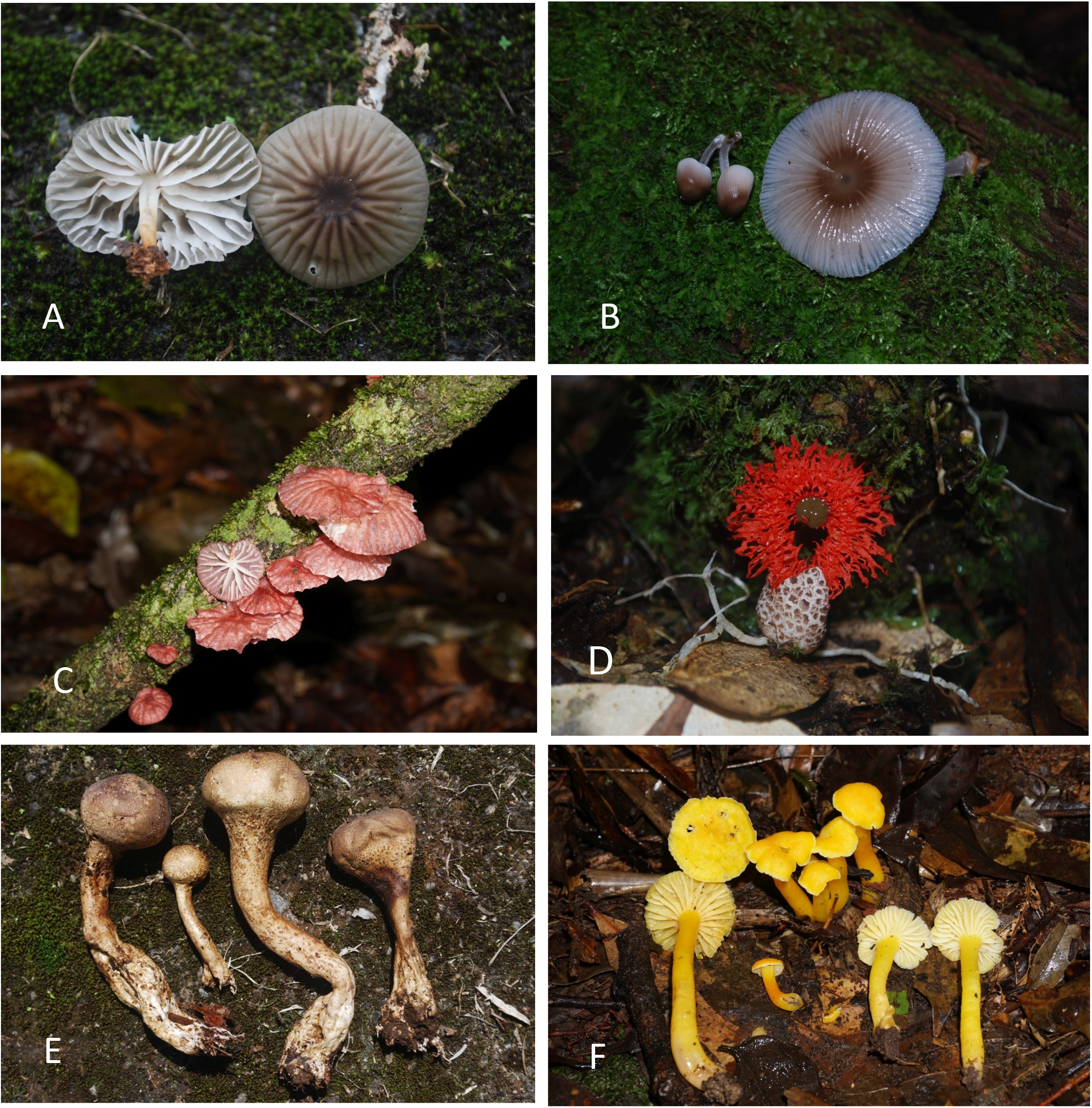
A-F. A. *Marasmius heliomyces*. B. *Mycena margarita*. C. *Crinipellis* sp. D. *Laternea pusilla.* E. *Veligaster nitidus*. F. *Hygrocybe* sp.

### *RUSSULA* DIVERSITY AT FORTUNA FOREST RESERVE

*Russula* has been found to be the most diverse genus of ECM fungi at Fortuna in both above and below ground surveys (Corrales 2016a, Corrales 2017). Results from our mushroom inventories show that *Russula* accounted for about 37% of the total number of ECM mushroom species and 25% of total number of species recorded at the study site including saprotrophs (Table 1). Molecular identification of fruiting body collections using Sanger sequencing indicated the existence of 40 OTUs, based on 97% similarity over the whole ITS region. This level of sequence similarity could be interpreted as consistent with species-level variation (Smith et al., 2007). The selected OTUs represented six *Russula* subgenera *sensu* Buyck et al. (2018). The most diverse subgenus was *Russula 2* with 19 OTUs, followed by subgenus *Heterophylla* with 9 OTUs, subgenus *Compactae* and *Crassotunicata* with 5 OTUs each, subgenus *Russula 1* with two OTUs, and finally subgenus *Malodora* with one OTU. Preliminary phylogenetic placement of the OTUs suggests that most, if not all, *Russula* species from Fortuna are new to science but more work is required to confirm this finding (B. Buyck *pers comm*). Only one OTU confirmed by collections Corrales 182 and 626 matched an existing species (*R. nigricans* Fr.) based on morphology and 99% similarity of the ITS region. However, since this is a European species more sequencing is required to confirm this name.

A remarkable feature of the Fortuna *Russula* community is its high diversity, containing species belonging to six out of eight subgenera recognized by Buyck et al. (2018). The only subgenera as yet uncollected at Fortuna are *Archea* and *Brevipes*. Also remarkable is that the Fortuna dataset is particularly rich in species in the subgenus *Crassotunicata* given that there are only 3 species described for this subgenus worldwide (Buyck et al. 2018). At present there are five OTUs of this subgenus found at Fortuna that do not match any of the described species (S. Adamčík *pers comm*).

### BIOGEOGRAPHIC CONSIDERATIONS OF THE ECTOMYCORRHIZAL COMMUNITIES AT FORTUNA

Many of the ECM host tree species in montane forests in Central and South America are considered relict species with a northern hemisphere temperate origin and therefore their associated fungal communities show a strong affinity with the ones found in North America (Halling, 2001). Pollen fossil records show that *Oreomunnea* has been present in Panama since the Early Miocene (Graham, 1989; Herrera et al., 2014) while *Quercus* arrived probably much later (Kappelle, 2006). Ectomycorrhizal fungal species associated with these host trees might have a particular evolutionary history given the high degree of isolation and potential speciation that they might have undergone. Mueller and Halling (1995), based on the analysis of the four best known ECM fungal genera associated with *Quercus* forests, established that in Central and South America the ECM fungal communities associated with this genus show a high degree of endemism, with most species having limited geographic ranges.

Comparisons of the ECM macromycete community composition of Fortuna with other cloud forests is difficult at this time because of a lack of comparable studies in neotropical montane sites that include intensive multi-year collecting over lengthy periods. Our approach of inventorying all fruiting bodies of the ECM fungi has yielded a wealth of diversity that will require the aid of specialists to assist at arriving at species determinations. Comparisons with fungal inventories from other tropical countries is currently only possible only at the genus level. However, future species-level comparisons will yield the most valuable biogeographic information.

Nearly all of the ECM genera that are reported here (Table 1) are common genera that can be found from the neotropics to high latitudes in North America (Corrales et al. 2018) as long as their associated ECM tree hosts are present. The only genus found at Fortuna that is likely to have a mainly or exclusively tropical distribution is *Veligaster* (Guzmán et al., 2004; Kasuya and Guzmán, 2007).

As many as 70% of the ECM fungal species in a newly explored area may be undescribed (Henkel et al., 2012). At this point we can only comment on the distribution of a few species that have been identified at Fortuna (Table S1). *Amanita* species were abundant at the study area and we have confirmed identifications for three species: *A. garabitoana* Tulloss, Halling & Mueller, *A. flavoconia* var. *inquinata* Tulloss, Ovrebo & Halling, and *A. brunneolocularis* Tulloss, Ovrebo & Halling. *Amanita garabitoana* has been recorded from Honduras to Costa Rica (Tulloss et al., 2011), and *A. flavoconia* var. *inquinata* and *A. brunneolocularis* are reported from Colombia and Costa Rica (Tulloss, 2005). Tulloss (2005) reports these species as occurring with *Quercus,* but we are unsure if these species are associated with *Quercus* or *Oreomunnea* in our study area given that they occur in forests where both host taxa are abundant. *Cortinarius* species were also abundant at the study area and we have identified two species: *Cortinarius neotropicus* Harrower and *Cortinarius costaricensis* Ammirati, Halling & Garnica. *Cortinarius neotropicus* was described from Costa Rica and has also been reported from Colombia (Harrower et al., 2015) as occurring under *Quercus*. At Fortuna this species was also found growing in *Quercus* dominated forest. *Cortinarius costaricensis* was described from under *Quercus* based on a single basidiome from Costa Rica; we have now made several additional collections to add to its distribution and to add *Oreomunnea* as a new potential host. We have also confirmed the identity of *Phylloporus centroamericanus* previously known only from the oak forests of Costa Rica (Neves and Halling, 2010) and *Phylloporus caballeroi* Singer described from Argentina and also reported for Bolivia, Costa Rica and Panama growing under *Alnus acuminata* forest (Neves and Halling, 2010). In Panama this species has been previously collected in the Parque Internacional La Amistad, Chiriquí, by R. Halling (REH 7906, NYBG). *Phellodon niger* (Fr.) P. Karst. occurs at Fortuna and is also known from the temperate forests of North America. For boletes, *Leccinum talamancae* Halling, Gómez & Lannoy and *L. tablense* Halling & G.M. Mueller were described from Costa Rica from under *Quercus* and we have now collected them at Fortuna. Notably, the ECM genus *Hebeloma* reported for Panama based on the STRI fungi checklist, has not yet been encountered at Fortuna. Also, absent from the STRI check list, and yet to be collected at Fortuna or sequenced from roots, is the genus *Tricholoma,* known to occur in the oak forests of Colombia and Costa Rica (Halling and Mueller, 2005; Mueller et al., 2006).

The new species and genus records from Fortuna are biogeographically relevant since they fill a gap of ECM records between the montane forests of Costa Rica and Colombia. Several ECM host plants including *Quercus*, *Oreomunnea*, and *Alfaroa* reach their most southern latitudinal distribution in Colombia and the Panama isthmus must have been an important bridge for migration of these species and their associated fungi. Research on the occurrence of fungi in Panama therefore provides important insights into the factors that control the diversity and biogeographical patterns of fungi in northern South America. The forests at Fortuna should continue to yield new ECM macromycete data in a number of ways that will help to improve our understanding of macromycete biodiversity and biogeography, including: (1) information on the fungal associates of *Oreomunnea mexicana* and *Quercus* spp., (2) data on the occurrence of fungi in the cloud forest ecosystem, (3) comparisons with the biodiversity of adjacent countries such as Colombia and Costa Rica where more field work has been done especially in the oak forests, (4) and contribution to the knowledge of fungal biodiversity of Panama. Finally, while ECM fungi have been the main focus of our work, saprotrophic macromycetes (and micromycetes) should not be neglected because they might be a source of important information on fungal biogeography and biodiversity especially compared to arbuscular mycorrhizal dominated forests where saprotrophs are generally the most abundant group of Agaricomycete fungi.

## CONCLUSIONS AND FUTURE DIRECTIONS

Based on our preliminary list of macrofungi in the Fortuna we conclude that this area harbors a high diversity of ECM fungi, and although not a particular focus of our work, there may be a rich diversity of saprotrophic fungi that future field work will reveal. Here we reported 22 new records of genera and 29 new records of species for Panama. In the near future we expect the number of species at Fortuna to grow considerably as we work closely with specialists to describe new species. These long-term taxonomic efforts will be important in improving the growing database of fungal records that is being compiled for Panama. Furthermore, exploring under-sampled ecosystems and new host species will improve the quality of reference voucher and DNA collections needed to inform environmental next generation sequencing methods.

We hope that our work continues to fill in the gap of knowledge regarding fungal diversity in montane cloud forests and also contributes to the database of knowledge of oak forest ECM fungi particularly in comparison to Costa Rica and Colombia where more intensive oak forest inventories have been made. Upon completion of the identification of our collections we should be able to make more accurate statements about the degree of endemism in the Fortuna region as well.

Given the functional importance of fungi in ecosystems and the high susceptibility of tropical montane forest to climate change, characterizing the fungal communities associated with this ecosystem and their variability along environmental gradients is essential for a better understanding of the microbial drivers of key ecosystems functions. Fungi play a key role in soil nutrient cycling and in the adaptation of plants to novel environmental conditions, two processes that are crucial under a global change scenery particularly in tropical montane forests.

## ACKNOWLEDGEMENTS

Funding from the Smithsonian Tropical Research Institute pre-doctoral Fellowship, NSF Dissertation Improvement Grant (Award Id, 1501483), COLCIENCIAS (497-2009), Robert L. Gilbertson Mycological Herbarium Grant (University of Arizona), Tinker Summer Research Fellowship, Francis M. and Harlie M. Clark Research support grant, and the Graduate College at University of Illinois are gratefully acknowledged. We thank Kayla Arendt, Jana U’Ren, Betsy Arnold, Katy Heath, and Pat Burke for assistance with molecular work and also Fredy Miranda, Jan Miranda, Evidelio Garcia, Carmen Velasquez, Carlos Espinosa, Marggie Rodriguez, Katie Heineman, Carlos Espinosa, and Cecilia Prada, and ENEL green power for help with fieldwork and logistic support. Finally, we thank Jim Dalling and Ben Turner for their help throughout the development of this project and to Meike Piepenbring for her helpful comments on a previous draft of the chapter.

We would also like to thank all the specialists that have helped us to identify the species from Fortuna: Michael A. Castellano (US Forest Service), Andrew Wilson (Denver Botanic Gardens), Gregory Mueller (The Field Museum), Brandon Matheny and Rachel Sweenie (UTK), Slavomir Adamčík (Slovak Academy of Science), Bart Buyck (Muséum national d’histoire naturelle, Paris), Donald H. Pfister (Harvard University), Roberto Garibay-Orijel (UNAM) and Leticia Montoya (INECOL), Ed Lickey (Bridgewater College), Roy Halling (NYBG), Joe Ammirati (University of Washington), Kare Liimatainen and Tuula Niskanen (Kew Royal Botanic Gardens). Special thanks to Slavomir Adamčík for his input on the *Russula* taxonomy section of this chapter.

## REFERENCES

Buyck, B., Kauff, F., Eyssartier, G., Couloux, A., Hofstetter, V., 2014. A multilocus phylogeny for worldwide *Cantharellus* (Cantharellales, Agaricomycetidae). Fungal Divers. 64, 101–121. https://doi.org/10.1007/s13225-013-0272-3

Buyck, B., Zoller, S., Hofstetter, V., 2018. Walking the thin line… ten years later: the dilemma of above- versus below-ground features to support phylogenies in the Russulaceae (Basidiomycota). Fungal Divers. 89, 267–292. https://doi.org/10.1007/s13225-018-0397-5

Corrales, A., Arnold, A.E.E., Ferrer, A., Turner, B.L.L., Dalling, J.W.W., 2016. Variation in ectomycorrhizal fungal communities associated with *Oreomunnea mexicana* (Juglandaceae) in a Neotropical montane forest. Mycorrhiza 26, 1–17. https://doi.org/10.1007/s00572-015-0641-8

Corrales, A., Henkel, T.W., Smith, M.E., 2018. Ectomycorrhizal associations in the tropics - biogeography, diversity patterns and ecosystem roles. New Phytol. https://doi.org/10.1111/nph.15151

Corrales, A., Turner, B.L.L., Tedersoo, L., Anslan, S., Dalling, J.W.W., 2017. Nitrogen addition alters ectomycorrhizal fungal communities and soil enzyme activities in a tropical montane forest. Fungal Ecol. 27, 14–23. https://doi.org/10.1016/j.funeco.2017.02.004

Corre, M.D., Veldkamp, E., Arnold, J., Wright, S.J., Corre, M.D., Veldkamp, E., Arnold, J., Wright, S.J., 2010. Impact of elevated N input on soil N cycling and losses in old-growth lowland and montane forests in Panama. Ecology 91, 1715–1729.

Del Olmo-Ruiz, M., García-Sandoval, R., Alcántara-Ayala, O., Véliz, M., Luna-Vega, I., Garcia-Sandoval, R., Alcantara-Ayala, O., Veliz, M., Luna-Vega, I., 2017. Current knowledge of fungi from Neotropical montane cloud forests: distributional patterns and composition. Biodivers. Conserv. 26, 1919–1942. https://doi.org/10.1007/s10531-017-1337-5

Ekblad, A., Wallander, H., Godbold, D.L., Cruz, C., Johnson, D., Baldrian, P., Björk, R.G., Epron, D., Kieliszewska-Rokicka, B., Kjøller, R., Kraigher, H., Matzner, E., Neumann, J., Plassard, C., 2013. The production and turnover of extramatrical mycelium of ectomycorrhizal fungi in forest soils: Role in carbon cycling. Plant Soil 366, 1–27. https://doi.org/10.1007/s11104-013-1630-3

Franco-Molano, A.E., Corrales, A., Vasco-Palacios, A.M., 2010. Macrofungi of Colombia II. Checklist of the species of Agaricales, Boletales, Cantharellales, and Russulales (Agaricomycetes, Basidiomycota). Actual. Biol. 32, 89–114.

Geml, J., Pastor, N., Fernandez, L., Pacheco, S., Semenova, T.A., Becerra, A.G., Wicaksono, C.Y., Nouhra, E.R., 2014. Large-scale fungal diversity assessment in the Andean Yungas forests reveals strong community turnover among forest types along an altitudinal gradient. Mol. Ecol. 23, 2452–2472. https://doi.org/10.1111/mec.12765

Gómez-Hernández, M., Williams-Linera, G., Guevara, R., Lodge, D.J., Gomez-Hernandez, M., Williams-Linera, G., Guevara, R., Lodge, D.J., 2012. Patterns of macromycete community assemblage along an elevation gradient: options for fungal gradient and metacommunity analyse. Biodivers. Conserv. 21, 2247–2268. https://doi.org/10.1007/s10531-011-0180-3

Graham, A., 1989. Studies in Neotropical paleobotany. VII. The Lower Miocene communities of Panama: The La Boca Formation. Ann. Missouri Bot. Gard. 76, 50–66.

Guzmán, G., Ramírez-Guillén, F., Miller, O.K., Lodge, D.J., Baroni, T.J., 2004. *Scleroderma stellatum* versus *Scleroderma bermudense*: the status of *Scleroderma echinatum* and the first record of Veligaster nitidum from the Virgin Islands. Mycologia 96, 1370–1379.

Halling, R.E., 2001. Ectomycorrhizae: Co-Evolution, Significance, and Biogeography. Ann. Missouri Bot. Gard. 88, 5–13.

Halling, R.E., Mueller, G.M., 2005. Common mushrooms of the Talamanca mountains, Costa Rica. The New York Botanical Garden, New York Bot Garden, Bronx, NY 10458 USA.

Halling, R.E., Mueller, G.M., 2002. Agarics and Boletes of Neotropical Oakwoods, in: Watling, R., Frankland, J.C., Ainsworth, A.M., Isaac, S., Robinson, C.H. (Eds.), Tropical Mycology, Vol 1. CABI Publishing, New York, pp. 1–10.

Halling, R.E., Mueller, G.M., 1999. New Boletes from Costa Rica. Mycologia 91, 893–899. https://doi.org/Doi10.2307/3761543

Halling, R.E., Ovrebo, C.L., 1987. A New Species of Rozites from Oak Forests of Colombia, with Notes on Biogeography. Mycologia 79, 674. https://doi.org/10.2307/3807818

Harrower, E., Bougher, N.L., Winterbottom, C., Henkel, T.W., Horak, E., Matheny, P.B., 2015. New species in *Cortinarius* section *Cortinarius* (Agaricales) from the Americas and Australasia. MycoKeys 11, 1–21. https://doi.org/10.3897/mycokeys.11.5409

Henkel, T.W., Aime, M.C., Chin, M.M.L., Miller, S.L., Vilgalys, R., Smith, M.E., 2012. Ectomycorrhizal fungal sporocarp diversity and discovery of new taxa in *Dicymbe* monodominant forests of the Guiana Shield. Biodivers. Conserv. 21, 2195–2220. https://doi.org/10.1007/s10531-011-0166-1

Herrera, F., Manchester, S.R., Koll, R., Jaramillo, C., 2014. Fruits of Oreomunnea (Juglandaceae) in the early Miocene of Panama. Paleobotany Biogeogr. A Festschrift Alan Graham his 80th year 124–133.

Kappelle, M., 2006. Ecology and conservation of Neotropical Montane Oak Forests. Ecol. Stud. 185.

Kasuya, T., Guzmán, G., 2007. *Veligaster nitidum*, a pantropical sclerodermataceous fungus new to Japan and Thailand. Mycoscience 48, 259–262. https://doi.org/10.1007/s10267-007-0359-3

Kennedy, P.G., Garibay-Orijel, R., Higgins, L.M., Angeles-Arguiz, R., 2011. Ectomycorrhizal fungi in Mexican *Alnus* forests support the host co-migration hypothesis and continental-scale patterns in phylogeography. Mycorrhiza 21, 559–568. https://doi.org/10.1007/s00572-011-0366-2

Kennedy, P.G., Walker, J.K.M., Bogar, L.M., 2015. Interspecific mycorrhizal networks and non-networking hosts: exploring the ecology of the host genus *Alnus*, in: Horton, T.R. (Ed.), Mycorrhizal Networks, Ecological Studies 224. Springer Science, Dordrecht, Netherlands, pp. 227–254.

Koljalg, U., Dahlberg, A., Taylor, A.F.S., Larsson, E., Hallenberg, N., Stenlid, J., Larsson, K.-H., Fransson, P.M., Karen, O., Jonsson, L., 2000. Diversity and abundance of resupinate thelephoroid fungi as ectomycorrhizal symbionts in Swedish boreal forests. Mol. Ecol. 9, 1985–1996.

Lilleskov, E.A., Hobbie, E.A., Horton, T.R., 2011. Conservation of ectomycorrhizal fungi: Exploring the linkages between functional and taxonomic responses to anthropogenic N deposition. Fungal Ecol. 4, 174–183. https://doi.org/10.1016/j.funeco.2010.09.008

Morris, M.H., Perez-Perez, M.A., Smith, M.E., Bledsoe, C.S., 2008. Multiple species of ectomycorrhizal fungi are frequently detected on individual oak root tips in a tropical cloud forest. Mycorrhiza 18, 375–383. https://doi.org/10.1007/s00572-008-0186-1

Mueller, G.M., Halling, R.E., 1995. Evidence for High Biodiversity of Agaricales (Fungi) in Neotropical Montane Quercus Forests. Biodivers. Conserv. Neotrop. Mont. For. 303–312.

Mueller, G.M., Halling, R.E., Carranza, J., Mata, M., Schmit, J.P., 2006. Saprotrophic and Ectomycorrhizal Macrofungi of Costa Rican Oak Forests, in: Kappelle, M. (Ed.), Ecology and Conservation of Neotropical Montane Oak Forests. Springer-Verlag Berlin Heidelberg, pp. 55–68.

Neves, M.A., Halling, R.E., 2010. Study on species of *phylloporus* I: Neotropics and North America. Mycologia 102, 923–943. https://doi.org/10.3852/09-215

Piepenbring, M., 2007. Inventoring the fungi of Panama. Biodivers. Conserv. 16, 73–84. https://doi.org/10.1007/s10531-006-9051-8

Piepenbring, M., 2006. Checklist of fungi in Panama, preliminary version. Nat. Rev. Cient ́ıfica y Hum. ́ıstica la Univ. Auto ́ noma Chiriquí 11.

Piepenbring, M., Hofmann, T.A., Miranda, E., Cáceres, O., Unterseher, M., 2015. Leaf shedding and weather in tropical dry-seasonal forest shape the phenology of fungi - Lessons from two years of monthly surveys in southwestern Panama. Fungal Ecol. 18, 83–92. https://doi.org/10.1016/j.funeco.2015.08.004

Põlme, S., Bahram, M., Yamanaka, T., Nara, K., Dai, Y.C., Grebenc, T., Kraigher, H., Toivonen, M., Wang, P.-H., Matsuda, Y., Naadel, T., Kennedy, P.G., Kõljalg, U., Tedersoo, L., 2013. Biogeography of ectomycorrhizal fungi associated with alders (*Alnus* spp.) in relation to biotic and abiotic variables at the global scale. New Phytol. 198, 1239–1249. https://doi.org/10.1111/nph.12170

Smith, M.E., Douhan, G.W., Rizzo, D.M., 2007. Intra-specific and intra-sporocarp ITS variation of ectomycorrhizal fungi as assessed by rDNA sequencing of sporocarps and pooled ectomycorrhizal roots from a Quercus woodland. Mycorrhiza 18, 15–22. https://doi.org/10.1007/s00572-007-0148-z

Tedersoo, L., Bahram, M., Polme, S., Koljalg, U., Yorou, N.S., Wijesundera, R., Ruiz, L. V., Vasco-Palacios, A.M., Thu, P.Q., Suija, A., Smith, M.E., Sharp, C., Saluveer, E., Saitta, A., Rosas, M., Riit, T., Ratkowsky, D., Pritsch, K., Poldmaa, K., Piepenbring, M., Phosri, C., Peterson, M., Parts, K., Partel, K., Otsing, E., Nouhra, E., Njouonkou, A.L., Nilsson, R.H., Morgado, L.N., Mayor, J., May, T.W., Majuakim, L., Lodge, D.J., Lee, S.S., Larsson, K.- H.H., Kohout, P., Hosaka, K., Hiiesalu, I., Henkel, T.W., Harend, H., Guo, L.-d. D., Greslebin, A., Grelet, G., Geml, J., Gates, G., Dunstan, W., Dunk, C., Drenkhan, R., Dearnaley, J., De Kesel, A., Dang, T., Chen, X., Buegger, F., Brearley, F.Q., Bonito, G., Anslan, S., Abell, S., Abarenkov, K., 2014. Global diversity and geography of soil fungi. Science (80-.). 346, 1256688–1256688. https://doi.org/10.1126/science.1256688

Tulloss, R.E., 2005. Amanita - distribution in the Americas with comparison to eastern and southern Asia and notes on spore character variation with latitude and ecology. Mycotaxon 93, 189–231.

Tulloss, R.E., Halling, R.E., Mueller, G.M., 2011. Studies in *Amanita* (Amanitaceae) of Central America. 1. Three new species from Costa Rica and Honduras. Mycotaxon 117, 165–205. https://doi.org/10.5248/117.165

van der Linde, S., Suz, L.M., Orme, C.D.L., Cox, F., Andreae, H., Asi, E., Atkinson, B., Benham, S., Carroll, C., Cools, N., De Vos, B., Dietrich, H.P., Eichhorn, J., Gehrmann, J., Grebenc, T., Gweon, H.S., Hansen, K., Jacob, F., Kristofel, F., Lech, P., Manninger, M., Martin, J., Meesenburg, H., Merila, P., Nicolas, M., Pavlenda, P., Rautio, P., Schaub, M., Schrock, H.W., Seidling, W., Šramek, V., Thimonier, A., Thomsen, I.M., Titeux, H., Vanguelova, E., Verstraeten, A., Vesterdal, L., Waldner, P., Wijk, S., Zhang, Y., Žlindra, D., Bidartondo, M.I., 2018. Environment and host as large-scale controls of ectomycorrhizal fungi. Nature 561, E42. https://doi.org/10.1038/s41586-018-0312-y

Wicaksono, C.Y., Aguirre-Guiterrez, J., Nouhra, E., Pastor, N., Raes, N., Pacheco, S., Geml, J., 2017. Contracting montane cloud forests: a case study of the Andean alder (*Alnusacuminata*) and associated fungi in the Yungas. Biotropica 49, 141–152. https://doi.org/10.1111/btp.12394

